# Disentangling covariate effects on single cell-resolved epigenomes with DeepDive

**DOI:** 10.1101/2025.09.30.679466

**Authors:** Andreas Fønss Møller, Jesper Grud Skat Madsen

**Affiliations:** Department of Biochemistry and Molecular Biology, University of Southern Denmark, Odense M, Denmark; The Novo Nordisk Foundation Center for Genomic Mechanisms of Disease, Broad Institute of MIT and Harvard, Cambridge, MA 02142, USA

## Abstract

Understanding the effects of individual biological factors from single cell-resolved epigenomic data is hindered by multicollinearity, particularly in human cohorts. We introduce DeepDive, a novel deep learning framework designed to systematically disentangle known and unknown sources of variation in single-nucleus ATAC-seq data. DeepDive accurately reconstructs chromatin accessibility, outperforms state-of-the-art methods with incomplete covariate information, and robustly recovers true biological signals from even highly entangled covariates, unlocking counter-factual, *what-if*, analyses. Applying DeepDive to pancreatic islet cells, we perform counter-factual analyses to prioritize covariates associated with a type 2 diabetes-linked beta cell subtype and nominate transcription regulators. DeepDive offers a powerful and unbiased tool for mechanistic discovery in complex human disease cohorts.

## Main

The epigenome of a single cell is a fingerprint of its identity and a record of its exposure to environmental, developmental, and genetic factors that, via dynamic and complex interactions, shape cellular function and fate through transcriptional regulation. We can map accessible chromatin in single nuclei at scale using single-nucleus Assay of Transposase Accessible Chromatin sequencing (snATAC-seq). Chromatin accessibility can be used for identification of cis-regulatory elements, and as a proxy for their activity. However, separating the effects of individual exposures on chromatin accessibility remains a hard problem, especially in human cohort studies where multicollinearity between exposures is common. Multicollinearity naturally occurs between related biological exposures; for example, it is well-established that obesity is a strong risk factor for developing type 2 diabetes. Thus, among type 2 diabetics, the prevalence of obesity is higher than among non-diabetics. Using conventional statistical approaches, such as those implemented in widely adopted toolkits for snATAC-seq analysis, including Signac^1^ and ArchR^2^, multicollinearity leads to poor and unstable estimates of the effect of each exposure. As a result, key biological signals can be obscured, complicating the interpretation of disease mechanisms and cellular states.

To address this challenge, we introduce DeepDive, a novel computational framework for semi-supervised probabilistic modelling of sources of variation in single cell-resolved epigenomic data for precise estimation of the effect of individual exposures, even among highly multicollinear factors. DeepDive is designed to be user-friendly. It is simple to train with just a few lines of code and has thorough documentation with guidance for training and interpretation. DeepDive consists of two networks (Figure 1A). The first network learns disentangled representations of known covariates using adversarial learning. The second network models the residual variation, which cannot be explained by known covariates, using a conditional variational autoencoder. This dual network configuration separates known and unknown sources of variation and, therefore, enables the quantification of unknown variance and the identification of latent covariates. Utilizing a multi-decoder framework^3^, DeepDive quantifies uncertainty in its own estimates, which can be directly used for differential accessibility testing. The model is trained end-to-end using a training scheme to maximise the information learned from known covariates. The trained DeepDive model accurately reconstructs chromatin accessibility in the human liver^4^ (Figure 1B, Supplemental Figure 1A-F).

**Figure 1:**
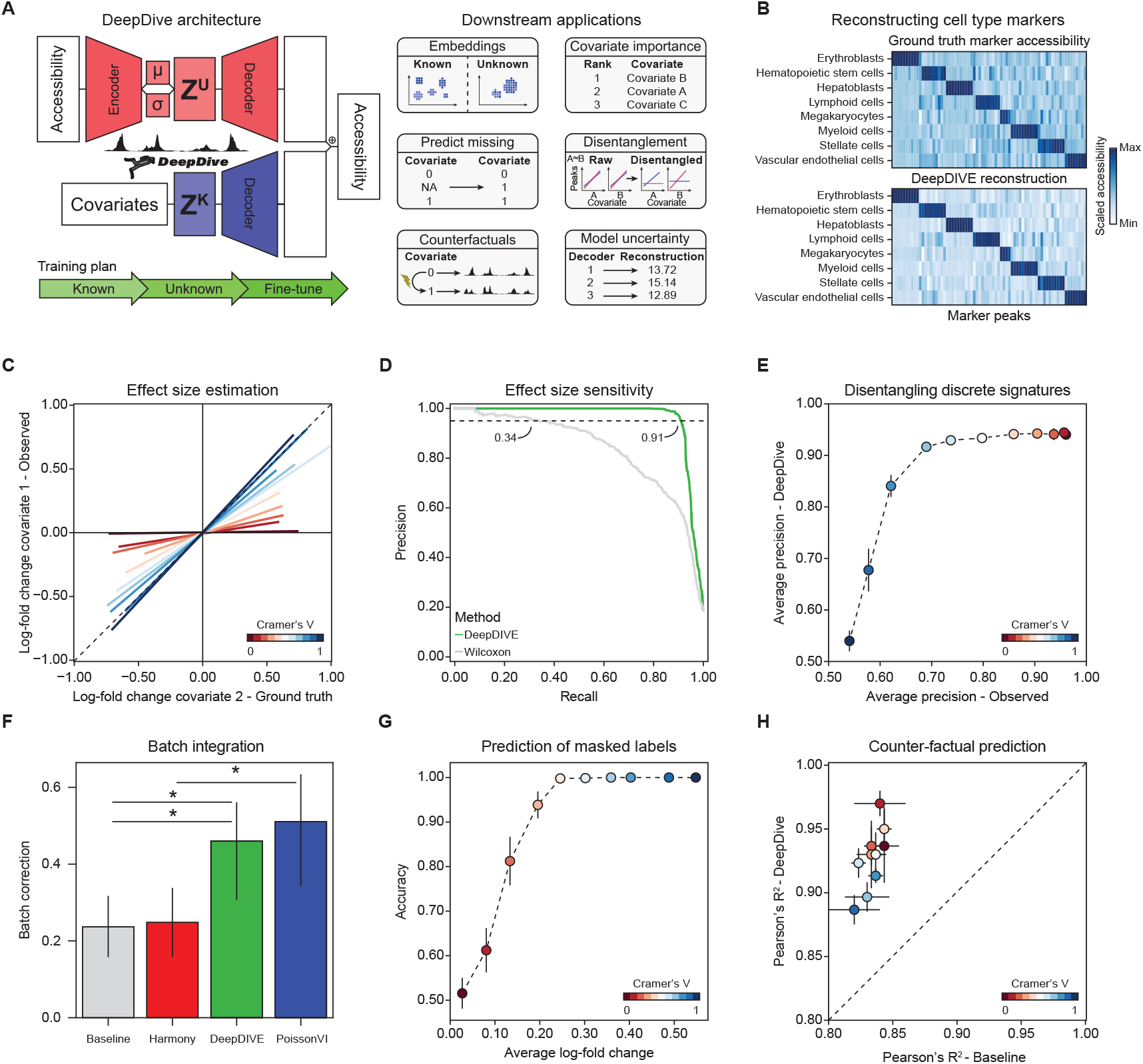
DeepDive accurately disentangles covariate effects on single cell epigenomes. **A)** Schematic overview of the DeepDive model. The model consists of a dual network architecture that disentangles known covariates using adversarial learning while modeling residual, unknown variation with a conditional variational autoencoder. **B)** Pseudobulk cell type marker peak chromatin accessibility of the raw dataset^4^ (top) and the DeepDive reconstruction after disentangling and removing donor-level covariates (bottom). **C)** Signal leakage from a simulated covariate to another as a function of increasing multicollinearity strength. **D)** Precision-recall curve comparing DeepDive’s and Wilcoxon Rank-sum’s performance in classifying differentially accessible peaks. **E)** Average precision for classifying differentially accessible peaks with discrete covariates, comparing DeepDive to log fold change-based classification, as a function of increasing multicollinearity strength. **F)** Average batch correction metric across the five datasets. Error bars represent the 95% confidence interval. **G)** Average accuracy of DeepDive’s predictions for masked covariate labels across effect sizes. **H)** Average Pearson’s R^2^ on hold-out covariate combinations.

In large-scale human studies, such as single cell atlas generation, where datasets are integrated prospectively, covariates describing important biological exposures may be partially or completely missing. For example, in the human lung cell atlas (HLCA), sex is missing for 47.5% of donors, age for 96.1% of donors, and smoking status for 32.6% of donors, and in the human retina cell atlas (HRCA), cause of death is missing for 80.8% of donors, while sex is missing for 7.7% of donors. To evaluate how imperfect information affects DeepDive, we trained DeepDive and BioLORD, a state-of-the-art model for covariate disentanglement of covariates^5^, on five different datasets with and without cell type labels. We omitted cell type labels because it is a strong covariate and, therefore, should be easy to separate in the basal latent space of each model, which captures residual variance. With cell type labels, DeepDive and BioLORD perform comparably, but in the absence of cell type labels, DeepDive is significantly more accurate (Supplemental Figure 2A-B) as it learns to separate cell types in basal latent space (Supplemental Figure 2C).

In addition to imperfect information, single cell atlases also often suffer from multicollinearity. For example, in the HLCA, there are varying degrees of multicollinearity with smoking status as cause of death is strongly collinear (Cramer’s V = 0.74) while sex is mildly collinear (Cramer’s V = 0.15). In the HRCA, there is also multicollinearity; for example, gender and age are collinear (Cramer’s V = 0.70). To benchmark how well DeepDive can disentangle the effects of collinear covariates, we initially simulated datasets with various levels of collinearity with a known ground truth (Supplemental Figure 3A-B). We confirmed that collinearity leads to poor effect size estimates as the biological signal from one covariate “leaks” to the other (Figure 1C), obscuring the true effects. At a moderate level of collinearity (Cramer’s V = 0.5), DeepDive accurately identifies 91% of differential features with an accepted error rate of 5%. In contrast, the Wilcoxon rank-sum test only identifies 34% (Figure 1D, Supplemental Figure 3C-E). Across all levels of collinearity, DeepDive consistently achieves similar or higher average precision for both discrete and continuous covariates (Figure 1E, Supplemental Figure 4A). Critically, this effect is dependent on our multi-decoder design (Supplemental Figure 4B-C) and is consistent across different levels of accessibility (Supplemental Figure 4D) and across different levels of missingness (Supplemental Figure 4E, Supplemental Table S1). To test the ability of DeepDive to disentangle covariates in real data, we subset a human islet of Langerhans dataset^6^ to introduce collinearity between sex and diabetes status and compared effect size estimates with the original dataset (Supplemental Figure 4F) and found that DeepDive accurately estimates effect sizes (Supplemental Figure 4G).

The adversarial training of the known covariate network in DeepDive encourages the disentangled representations of observed exposures to be independent. This design enables counter-factual prediction, where the model is tasked with predicting chromatin accessibility for unseen covariate combinations. To test the accuracy of counter-factual prediction with DeepDive, we devised three tasks inspired by potential use cases. In the first task, we use DeepDive for counter-factual prediction, omitting a covariate. We chose to omit the batch label, essentially using DeepDive for batch integration and evaluate across five datasets using scIB^7^. We found that DeepDive performs on par with contemporary state-of-the-art batch integration methods for snATAC-seq data (Figure 1F, Supplemental Figure 4H). In the second task, we use DeepDive for counter-factual prediction of missing donor covariates. To evaluate, we masked donor-level covariates with various effect sizes in simulated datasets and found that at moderate effect sizes, DeepDive could predict the missing covariate with high accuracy (Figure 1G). Similarly, we found that DeepDive significantly outperforms baseline models in predicting diabetes status in real data (Supplemental Figure 4I). In the third task, we use DeepDive to counter-factual predict chromatin accessibility of unseen combinations of covariates. To evaluate, we hold-out a covariate combination from simulated datasets and compare counter-factual prediction to the hold-out ground truth. As an illustrative example, in a dataset with two cell types (A and B) and two donors (X and Y), we hold-out cell type A in donor X, but A remains observed in donor Y. We evaluated DeepDive by virtually shifting the label of cell type B to A in donor X and comparing the predicted profile to the held-out data. We found that the counter-factual profile is significantly more similar to the ground truth than the baseline label (Figure 1H). To evaluate in real data, we held out a cell type in one donor and compared the counter-factual epigenome to the ground truth. We found that DeepDive accurately reconstructs the unseen epigenome, more accurately than using the average profile of the same cell type from other donors (Supplemental Figure 4J-K). In summary, DeepDive can disentangle collinear covariates for accurate reconstruction and counter-factual prediction of single cell-resolved epigenomes.

As a vignette of how counter-factual prediction with DeepDive can be used to nominate cell types and regulators linked to specific covariates, we trained a DeepDive model on snATAC-seq performed on human islets of Langerhans (Figure 2A) accounting for both biological covariates, including cell type, disease status, BMI, HbA1c, age, gender, and race, and technical covariates, including center, storage, purity, and fraction of mapped reads (Supplemental Methods). DeepDive accurately reconstructs the dataset from the known covariates, although these do not explain the entire epigenome (Supplemental Figure 5A). We initially focused on the interaction between obesity (as proxied by BMI) and diabetes status, which is known to be entangled^8^. To prioritize cell types affected by BMI and diabetes status, we used DeepDive to perform cell type-resolved covariate importance and found that both covariates most strongly affect beta cells (Figure 2B). To validate that DeepDive has disentangled these covariates, we identified differential accessibility peaks associated with each covariate and found the two peak sets to be orthogonal (Jaccard index = 0.1, Supplemental Figure 5B). Further, we leveraged human genetics and evaluated enrichment for genetic variants associated with type 2 diabetes and BMI. We found the peaks associated with BMI in DeepDive in beta cells are significantly enriched for variants linked to obesity, but not for variants linked to type 2 diabetes (Supplemental Figure 5C). Next, we set out to dissect how each donor-level covariate affects the proportion of the beta-1 and beta-2 beta cell subtypes, whose proportion has recently been shown to be strongly associated with HbA1c levels and diabetes status^6^. We reconstructed chromatin accessibility without technical confounders and trained an XGBoost model to classify beta-cells as beta-1 or beta-2 based on the authors original labels (Supplemental Figure 5D-F). Next, we virtually shifted donor covariates and found that disease status has the strongest effect on the proportion of the two states, followed by small additive effects from HbA1c, BMI, and age (Figure 2E) across both genders (Supplemental Figure 5G). To nominate transcription factors involved in the transition between these two beta-cell states, we used chromVAR^9^ to obtain motif activity estimates using the reconstructed chromatin accessibility profiles and using counter-factual profiles virtually shifting the diabetes status label (Supplemental Figure 5H). We identified 226 transcription factors associated with motifs differentially active between beta-1 and beta-2. Of those, the differential activity of 188 were rescued in counter-factual prediction (supported) and 38 were not (unsupported) (Supplemental Table S2). The regulators supported by counter-factual prediction were more enriched among proteins linked to type 2 diabetes through genetic evidence (Figure 5D) and correlated to glucose-stimulated insulin secretion (Figure 5E, Supplemental Figure 5I). This strongly indicates counter-factual prediction with DeepDive nominates high-confidence transcriptional regulators of significant interest for follow-up experiments.

**Figure 2:**
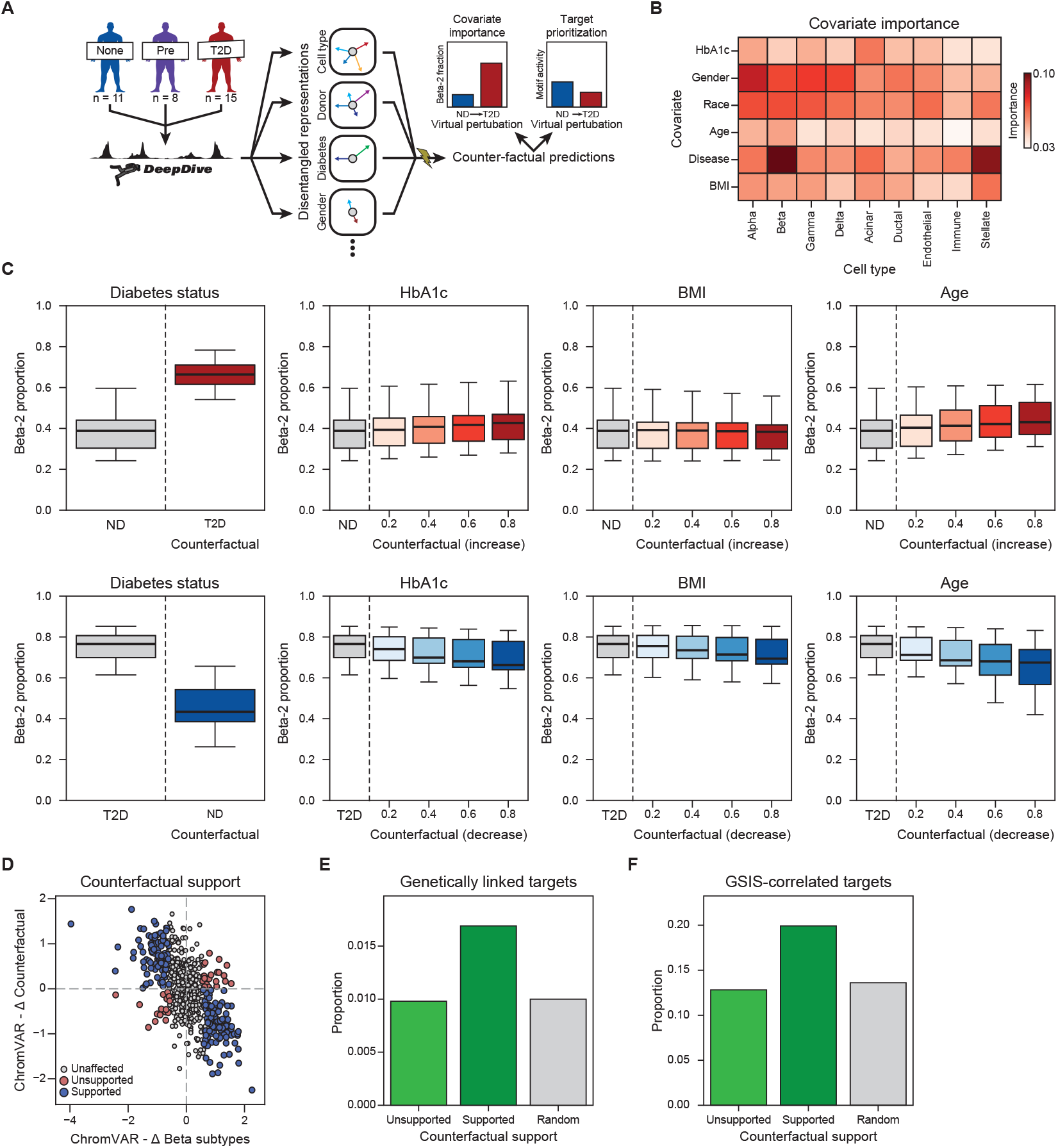
DeepDive disentangles covariate effects in pancreatic islet cells. **A)** Schematic overview. **B)** Cell type-specific covariate importance of the DeepDive model, showing the relative impact of each covariate on different cell types. **C)** Counterfactual prediction on non-diabetic (ND) (top panel) and type-2 diabetics (T2D) individuals (bottom panel), as measured by predicted beta-2 cell proportion. **E-F)** The proportion of transcription factors activities associated with beta-1 and beta-2 differences, stratified by DeepDive counterfactual support and random sampling, overlapping with eQTL and metabolic genome-wide association study (GWAS) colocalization data (E) or glucose-stimulated insulin secretion (GSIS) correlated targets (F).

DeepDive has limitations; DeepDive assumes that all covariates carry a signal, are independent, and linearly additive in latent space. We envision that future work can relax these limitations, for example, by using multiplicative, rather than additive, fusion, which would allow for non-linear interactions between covariates. In conclusion, we present DeepDive–an end-to-end probabilistic framework–capable of disentangling covariate effects on single cell-resolved epigenomes, thereby enabling accurate counter-factual prediction.

## Methods

### DeepDive

DeepDive is a deep generative model designed to disentangle known and unknown sources of variation in high-dimensional count data from single cell-resolved epigenomics assays. It combines a variational autoencoder (VAE) framework with adversarial networks to separate structured variation attributable to known covariates from residual, unannotated signals. The model specifics are described in greater detail in Supplemental Note 1.

In DeepDive, observed single cell-resolved profiles are modelled as arising from a latent representation that is partitioned into two components: (i) covariate-associated latent variables that capture structured effects of known sample-or cell-level annotations (e.g., donor phenotype or BMI), and (ii) residual latent variables that capture remaining biological or technical variation. Adversarial classifiers and regressors are trained jointly with the VAE to minimize the predictive power of covariates in the residual latent space, ensuring effective disentanglement of annotated and unannotated effects. DeepDive estimates model uncertainty using a multi-decoder setup^3^.

The generative model reconstructs high-dimensional accessibility profiles directly from these disentangled representations, enabling both counterfactual prediction and downstream interpretability. Specifically, by holding the residual latent space fixed while manipulating covariate-associated latent space, DeepDive can simulate chromatin accessibility under hypothetical covariate settings, thereby estimating covariate-specific effects.

### Training procedure

DeepDive was implemented with two hidden layers of 128 and 64 fully connected nodes in both the encoders and decoders, and a latent space of 32 dimensions. Adversarial networks were trained with a single hidden layer of 16 fully connected nodes. Unless otherwise indicates, DeepDive was trained using 20 decoders. During training, DeepDive randomly selects one decoder to update for every minibatch. Dropout was applied at a rate of 0.1 in the autoencoder components and 0.3 in the adversarial networks.

DeepDive was trained in three phases. In the first phase, only the covariate-associated (known) network was optimized. In the second phase, training was restricted to the residual (unknown) network while the known network weights were frozen. In the final phase, the known network was fine-tuned with the unknown network weights held fixed. The model was trained for 300 epochs per decoder, evenly distributed across the three phases. Optimization was performed using AdamW with learning rates of 0.0001 for autoencoders and 0.0003 for adversaries, and weight decay values of 0.001 and 0.0000004, respectively.

### Effect size estimation and differential analysis

To estimate differential effects attributable to a covariate, we compared decoder reconstructions under different covariate embeddings while holding all other background covariates constant. For discrete covariates, group-specific embeddings (e.g., group A vs. group B) were obtained from the model’s learned embedding space. For continuous covariates, the embedding was evaluated at the specified value and compared against a reference (zero). Where relevant, additional background covariates were fixed by adding their corresponding embeddings to the latent representation prior to reconstruction. For each covariate configuration, reconstructed feature values were obtained from all decoder networks. Feature-level differences between group A and group B reconstructions were computed per decoder. Within each decoder, a standardized test statistic was calculated by normalizing the difference across features to obtain a Z-score, and a corresponding two-sided P-value was derived from the standard normal distribution. To combine evidence across multiple decoders, P-values for each feature were aggregated using Fisher’s method:

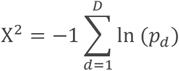

Where *D* is the number of decoders. Effect sizes were summarized as the mean reconstructed difference across decoders. Multiple testing correction was applied to combined *P*-values across all features using the Benjamini– Hochberg false discovery rate (FDR) procedure.

### Evaluation of effect size estimates

To evaluate the differential effect size estimation procedure, a simulation of single-nucleus chromatin accessibility profiles under controlled multicollinearity between discrete (Cramer’s V 0-1) and continuous (Pearson correlations 0-1) covariates was utilized. The presence of a non-zero simulated effect as the ground truth.

Estimates were likewise evaluated on beta cells from a real scATAC-seq data of pancreatic islets^6^ downsampled (20k cells) to induce multicollinearity between disease status and sex (Cramer’s V 0-1). Ground truth differentially accessible peaks were identified in rebalanced data (40k cells, Cramer’s V ∼0).

Performance was quantified using average precision from the precision-recall curve. DeepDive predicted effect sizes were compared to log-fold changes calculated on the simulated and downsampled data. Effect size estimation with log fold change and Wilcoxon Rank Sum differential accessibility testing was performed with scanpy^10^ rank_genes_groups with method = wilcoxon on log-transformed counts.

### Simulating count data with controllable covariate dependence

A simple simulation framework was implemented to generate single-nucleus chromatin accessibility profiles with controlled covariate influence. The simulation produces fragment counts with a specified number of cells and genomic features (peaks), while allowing covariates to modulate accessibility.

Each dataset *X* ∈ ℕ^*n*×m^ is initialized with baseline accessibility values drawn from a uniform distribution across features *μ*_*j*_*∼*Uniform(0.5, 2.0), *j* = 1, …, *m*. Cells are then assigned to covariate groups with probability p. If assuming dependency between two covariates, these are drawn from a joint probability distribution *P* ∈ ℝ^*r*×c^ specifying the relationship, where *r* and *c* are the number of groups in the two covariates. To get this distribution only parameterized by an entanglement score *η* ∈ [0, 1], where 0 is independence and 1 is perfect dependence, *P*(*η*) = (1 − *η*)*U* + *η*Ĩ, where U is a uniform joint distribution *U*_*ij*_ = 1/*rc* and Ĩ, = 1/ min(r, c) pad(*I*_min (*r,c*)_, thereby interpolating between perfect independence and dependence. Covariates are assumed to have independent effects, and thus, feature-level effect sizes are introduced by randomly selecting a subset of features for each covariate with proportion qc and applying multiplicative adjustments to their baseline accessibility. For each covariate *c* and group *g*, an effect vector is defined as 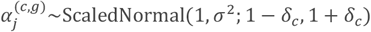 where *δ* is the maximum effect size applied to selected features and 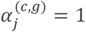 to non-selected features. For cell *i*, the mean accessibility of feature *j* is therefore given by 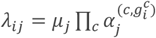. The final counts are finally sampled from a poisson distribution with mean values after adjustment *X*_*ij*_ *∼*Poisson(*λ*_*ij*_ ).

### Benchmarking of integration methods

Integration performance was benchmarked using the scIB framework^7^. We compared four methods: Latent Semantic Indexing (uncorrected), Harmony^11^, PoissonVI^12^, and DeepDive. The methods were applied to five publicly available scATAC-seq datasets^4,6,13-15^, spanning 27034–720613 cells and 5–34 donors.

Benchmarking was conducted using scIB^7^, which evaluates integration along two complementary axes: biological conservation and batch correction. Biological conservation was quantified using metrics of cell type separation and neighbourhood preservation, including normalized mutual information (NMI) and adjusted Rand index (ARI) with Leiden clustering, silhouette width with respect to cell type annotations, and the cell type–local inverse Simpson’s index (cLISI). Batch correction performance was assessed using the local inverse Simpson’s index (iLISI) for batch mixing, graph connectivity, and additional global measures of batch effect removal.

### Datasets and preprocessing

We used the following datasets and covariates:

1. NeuroIPS^14^ (GSE194122): Cell type, batch, site, DonorID, DonorBloodType, DonorRace, Ethnicity, DonorGender, DonorSmoker, DonorAge and DonorBMI.
2. Kidney^13^ (GSE151302): Celltype, sample. Age, gender, race and egfr.
3. GCC^15^ (GSE162170): Iterative.LSI.Clusters, Sample.ID, Age, Tissue.ID, Sample.Type, Batch and Age.
4. sciATAC-seq3^4^ (GSE149683): Cell_type, donor_id, sex, batch, tissue and day_of_pregnancy.
5. Pancreas^6^ (GSE169453): Cell_type, donor, Diabetes status, Human islet resource center, sex, ethnicity, 10x multiome assay, islet index, age, BMI, HbA1c and purity.

In benchmarking all datasets, peaks accessible in 10% of cells and cells with more than 5 accessible peaks were retained for analysis. In the vignette of disentangling the the effects to diabetes status and obesity, the thresholds were relaxed to peaks accessible in 1% of cells.

### Disentanglement with BioLORD

BioLORD^5^ was applied to the single cell-resolved accessibility data to learn disentangled representations of biological and technical covariates. The model was run following the authors’ recommended parameters as described in the official [vignette](https://biolord.readthedocs.io/en/latest/tutorials/biolord_omics.html).

### Counterfactual prediction of missing covariate combinations

To evaluate whether DeepDive can generalize to unseen combinations of covariates, a controlled simulation study was performed. Single cell-resolved chromatin accessibility profiles were simulated under two binary conditions with varying multicollinearity (Cramer’s V 0-1). To assess counterfactual prediction, all cells corresponding to one covariate combination were removed from the training data. A DeepDive model was trained on the remaining cells. The trained model was subsequently used to reconstruct chromatin accessibility profiles for: (i) cells from an observed covariate state, and (ii) a counterfactual prediction with a covariate switched to the unobserved covariate. Predicted accessibility profiles were compared against the true profiles that were held out and evaluated for non-zero effect-size features. Model accuracy was quantified by computing Pearson coefficient of determination (R2) between the mean predicted and ground truth accessibility.

### Predicting missing covariates

#### Cell-level covariates

DeepDive’s counterfactual reconstruction was used to impute missing cell-level covariates. For each covariate with missing values, we ranked its relative importance to define an inference order. Candidate values were taken as all observed categories for discrete covariates and as evenly spaced samples across the observed range for continuous covariates. Cells with missing values were reconstructed under each candidate while holding other covariates fixed. Reconstruction accuracy was assessed using the negative log-likelihood between reconstructed and observed accessibility profiles, and the candidate minimizing this error was assigned as the predicted covariate.

#### Donor-level covariates

For donor-level inference, we masked the diabetes status of a single donor and trained DeepDive while excluding donor-specific variation. We then generated non-diabetic (ND) and type 2 diabetic (T2D) counterfactual reconstructions for all unmasked donors. Differences between ND and T2D reconstructions, together with cell type labels, were used to train an XGBoost classifier. For the masked donor, predictions were made on its counterfactual differences, and cells with class probabilities ≥ 0.975 contributed to a majority vote used to assign the donor label.

### LD score regression

DeepDive was applied to identify peaks uniquely associated with BMI and T2D (FDR < 0.05, effect size > 0.1). Peaks were lifted over from hg38 to hg19 and overlapped with 1000 Genomes Project Phase 3 EUR SNPs within a 1,000 bp window centered on each peak. Partitioned heritability was estimated using stratified linkage disequilibrium score regression (S-LDSC)^16^ with the baselineLD v2.2 model.

### Counterfactual support

Motif activity analysis pyChromVAR^9^ was used to estimate motif activity on reconstructed profiles and T2D-to-ND counterfactuals, adjusting for the covariates cell type, disease, sex, race, age, BMI, and HbA1c, with default parameters. Motifs showing an absolute difference in chromVAR deviation > 1 between beta-1 and beta-2 cells were classified as regulated, while those below this threshold were considered unregulated. Regulated motifs were further stratified by direction of change in T2D-to-ND counterfactuals: motifs reversing the beta-subtype effect were designated “supported,” whereas those with the same directional effect were designated “unsupported.” Supported and unsupported motif sets were then tested for enrichment of transcription factors (TFs) associated with glucose-stimulated insulin secretion (GSIS) or with genetic evidence.

GSIS-associated TFs were defined by per-gene ordinary least squares regression, modelling the relationship between gene expression and the GSIS stimulation index while adjusting for culture time, donor age, donor sex, cold ischemia time, islet purity, cell trapping percentage, and digestion time (FDR < 0.01; expressed in ≥1% of pancreatic islet cells^6^).

TFs with genetic support were obtained from a database of eQTLs colocalizing with GWAS loci for metabolic traits^17^, restricted to TFs expressed in ≥1% of pancreatic islet cells^6^.

### Statistics and reproducibility

The statistical analysis of the data was described in the above methods and can be reproduced from scripts available on GitHub (http://github.com/madsen-lab/DeepDive_reproducibility). No statistical tests were used to predetermine sample size. When simulating or downsampling datasets, the process was repeated three times, and average scores were reported. The investigators were not blinded to allocation during experiments and outcome assessment.

## Data availability

All datasets analyzed or used for benchmarking are publicly available and summarized in the previous data processing section.

## Code availability

DeepDive is available as a Python package at http://github.com/madsen-lab/DeepDive. The code to reproduce the results is available at http://github.com/madsen-lab/DeepDive_reproducibility.

## Author contribution

A.F.M. and J.G.S.M. conceptualized the method. A.F.M and J.G.S.M implemented the code, performed analysis, and prepared figures. J.G.S.M. supervised the work. J.G.S.M. wrote the manuscript with inputs from A.F.M. All authors contributed to its editing.

## Acknowledgments

This work was supported by grants from the Novo Nordisk Foundation (NNF21SA0072102, J.G.S.M. and NNF21OC0068929, J.G.S.M.), as well as the Danish National Research Foundation (DNRF141, J.G.S.M.) to the Center for Functional Genomics and Tissue Plasticity (ATLAS). Computation for this project was performed using the UCloud interactive HPC system, which is managed by the eScience Center at the University of Southern Denmark.

