## Supplemental Figure for "Disentangling covariate effects on single cell-resolved epigenomes with DeepDive"

### Supplemental Figure 1

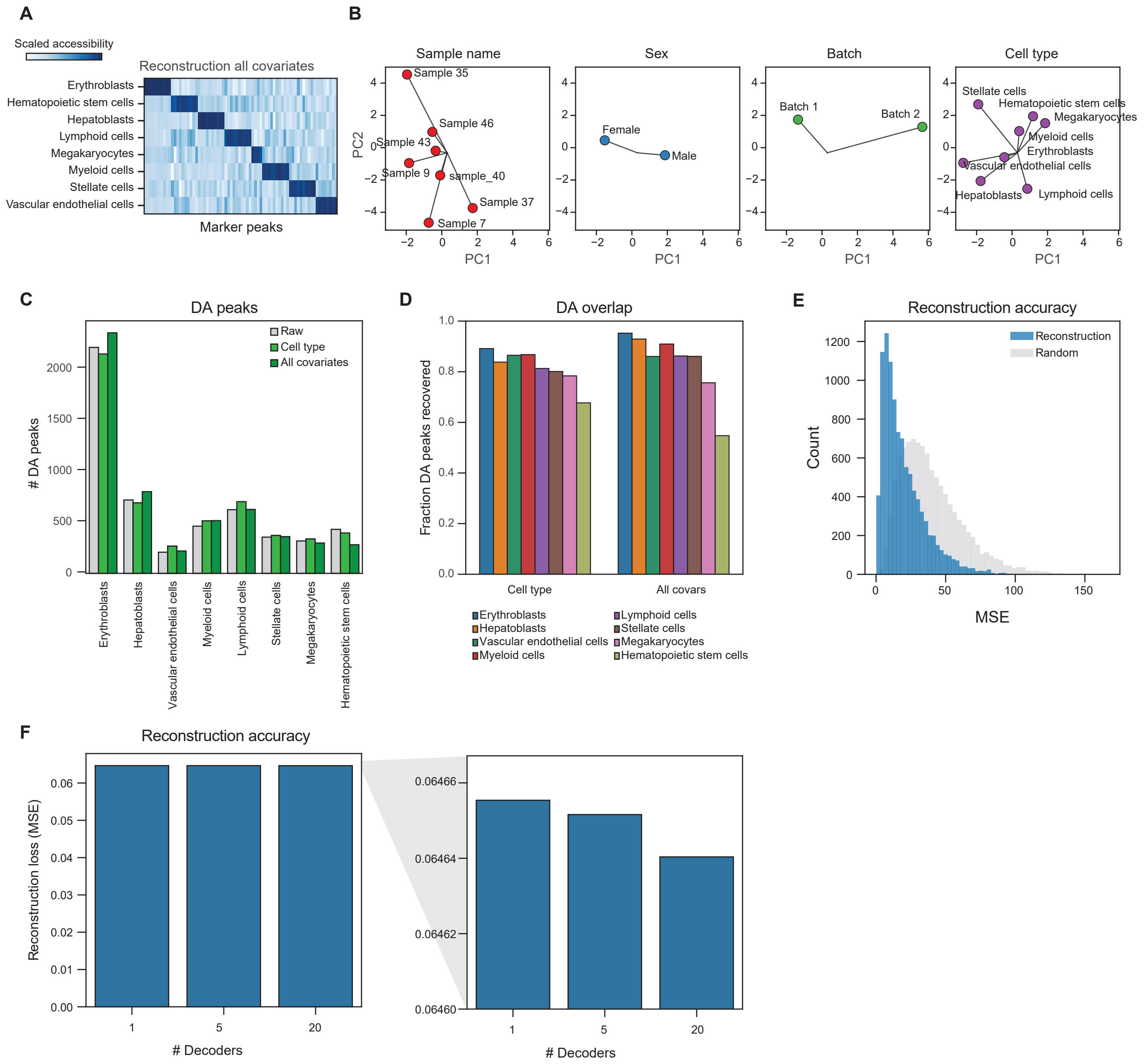

### Supplemental Figure 2

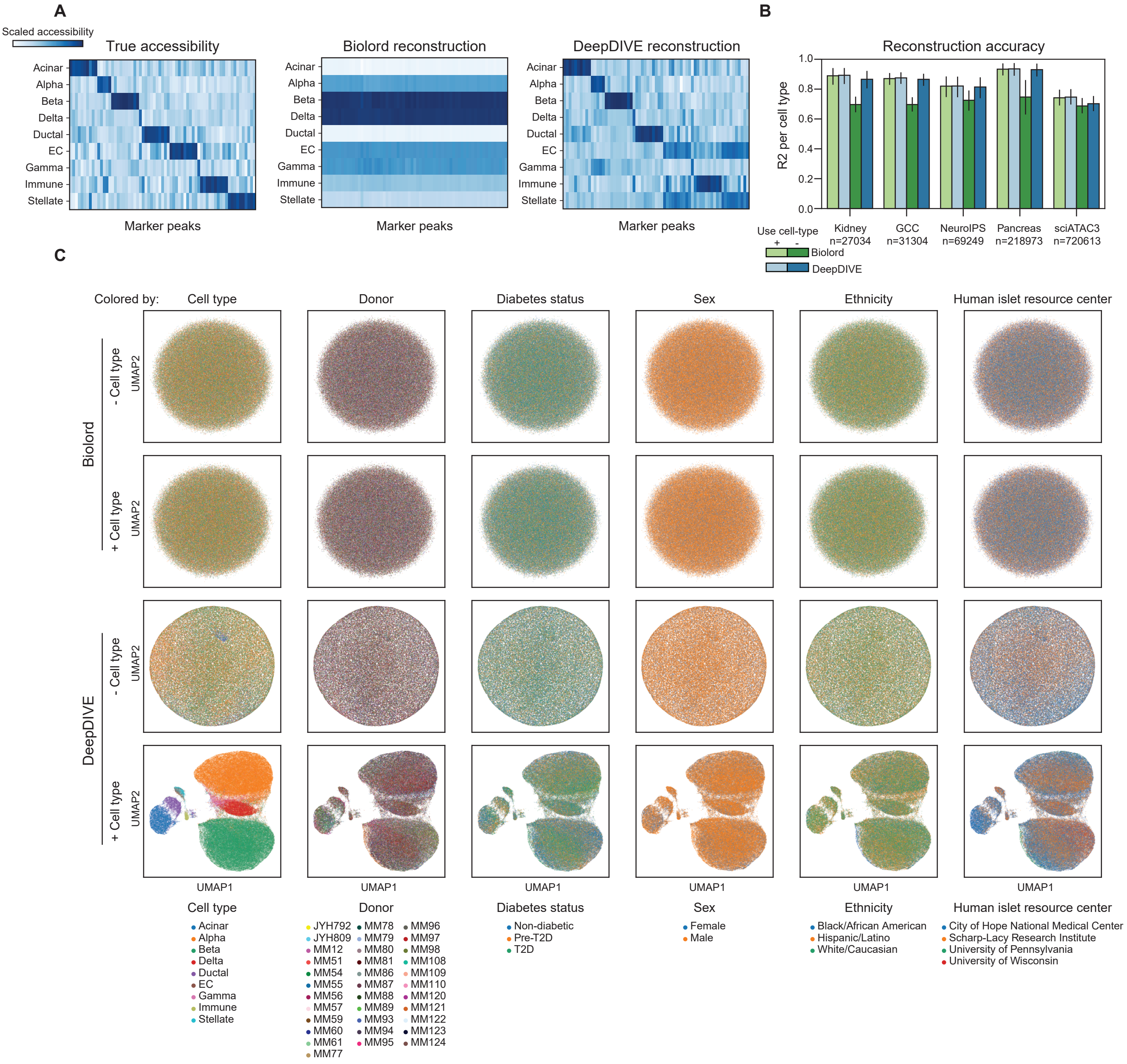

### Supplemental Figure 3

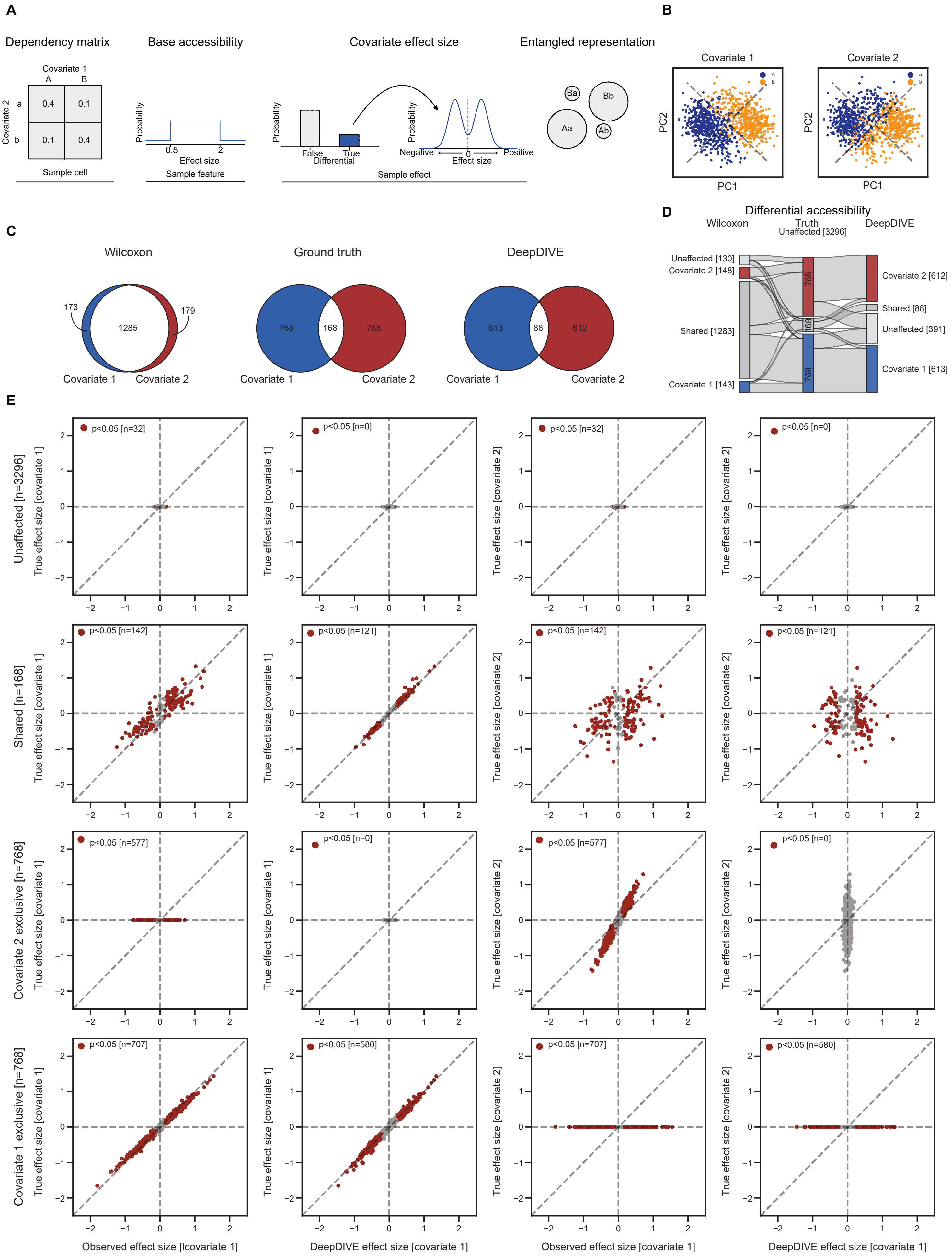

### Supplemental Figure 4

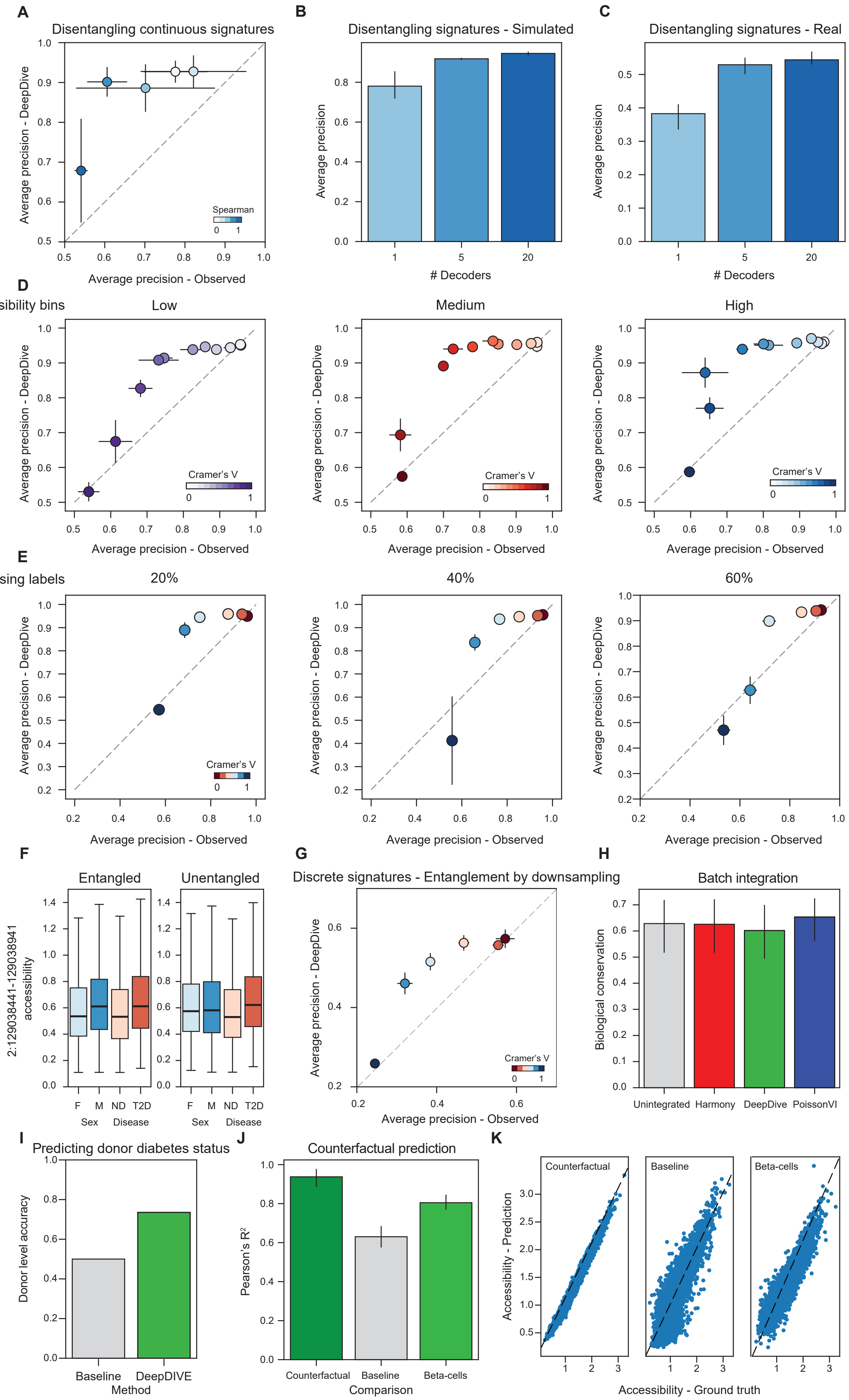

### Supplemental Figure 5

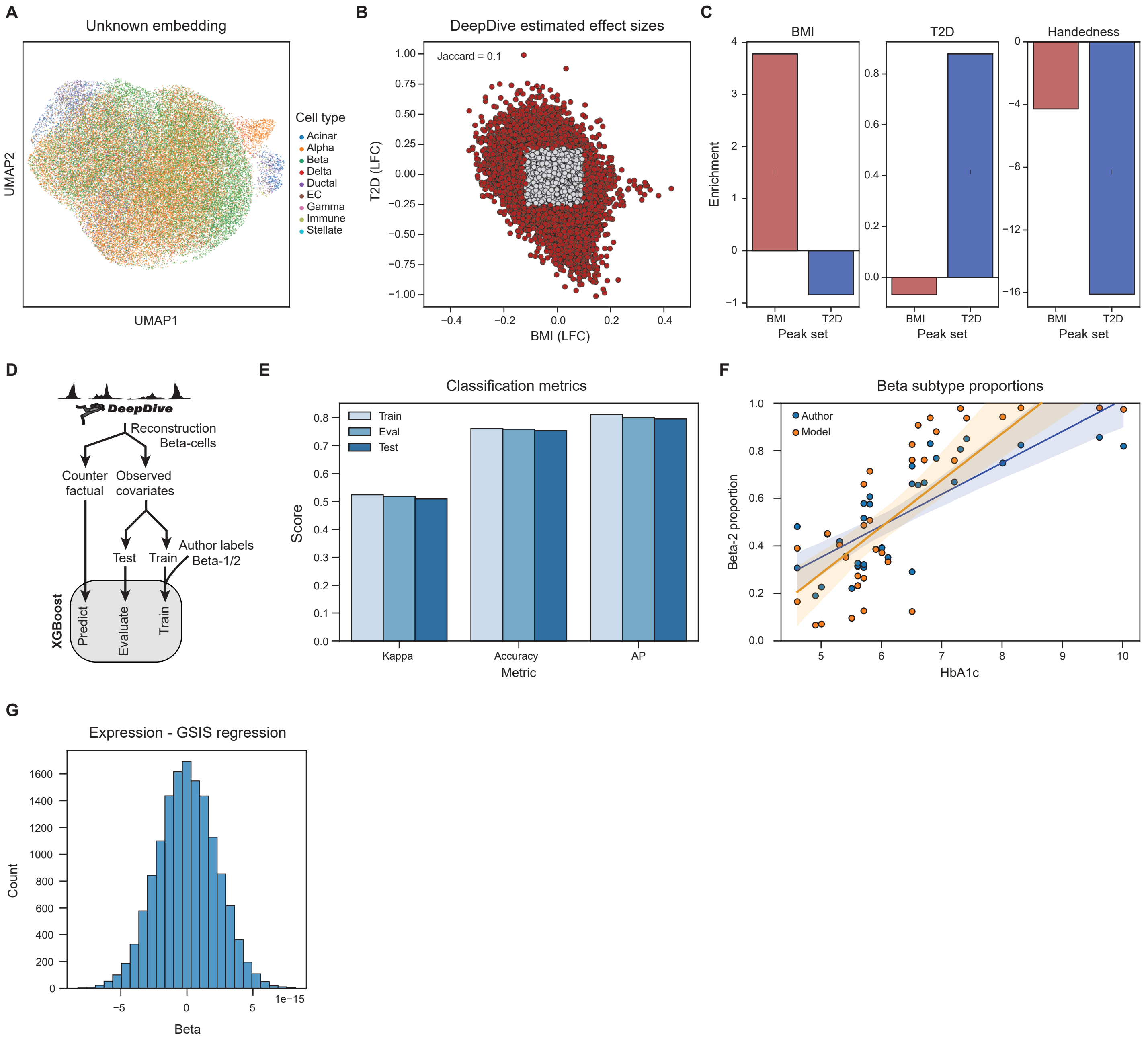

#### Supplemental Figure legends

**Supplementary Figure 1:**
**A)** Pseudobulk chromatin accessibility of the DeepDive reconstruction using all covariates[4]. **B)** PCA of covariate embeddings for known covariates, illustrating how DeepDive represents and disentangles these factors in the latent space. **C)** Number of marker peaks identified from the raw data and the DeepDive reconstructions. **D)** Fraction of marker peaks identified from the raw data that are successfully recovered in the DeepDive reconstruction. Reconstruction with only cell type labels on the left and reconstruction with all covariates on the right. **E)** Per cell mean squared error (MSE) of the reconstruction and between random cell pairs. **F)** MSE of reconstruction on DeepDive models trained with 1, 5, and 20 decoders.

**Supplementary Figure 2:**
**A)** Accessibility of marker peaks from the human pancreas raw dataset (left) compared to reconstructions by Biolord (middle) and DeepDive (right). Both models were trained without explicit cell type information. **B)** Average reconstruction accuracy ($R^2$) of cell type pseudobulk profiles, comparing Biolord and DeepDive models trained both with and without cell type information, across the five benchmarking datasets. Error bars represent the 95% confidence interval. **C)** UMAP visualizations of basal embeddings from the pancreas dataset[5], generated by Biolord (top two rows) and DeepDive (bottom two rows). The top row for each model shows training with cell type information, while the bottom row shows training without.

**Supplementary Figure 3:**
**A)** Schematic illustrating the process of simulating scATAC-seq data with controlled multicollinearity between covariates. **B)** UMAP visualizations of the simulated dataset, displaying the distributions according to two entangled covariates with a Cramer's V of 0.5. **C)** Venn diagrams comparing simulated differentially accessible peaks (middle) with differential peaks classified using Wilcoxon Rank-Sum (left) and DeepDive (right) under multicollinear conditions. **D)** Sankey plot visualizing the alignment between peak classification by Wilcoxon Rank-Sum and DeepDive, and the ground truth differentially accessible peaks. **E)** Accuracy of the estimated effect size for two entangled covariates, comparing estimates derived from log fold change calculations and hypothesis testing using Wilcoxon Rank-Sum versus DeepDive.

**Supplementary Figure 4:**
**A)** Average precision for classifying differentially accessible peaks with continuous covariates, comparing DeepDive to log fold change-based classification (by grouping covariate values as high and low), as a function of increasing multicollinearity strength. **B)** Average precision for classifying differentially accessible (DA) peaks at a Cramer's V of 0.6 with 1, 5, and 20 decoders, in simulated data. **C)** Average precision for classifying DA peaks at a Cramer's V of 0.6 between disease status and sex, with 1, 5, and 20 decoders, achieved by downsampling the pancreas dataset[5]. **D-E)** DeepDive maintains high average precision when classifying differentially accessible peaks with discrete covariates across different accessibility bins (low, medium, and high) (D) and levels of missingness (20%, 40% and 60%) (E). **F-G)** Classifying differentially accessible peaks with discrete covariates, comparing DeepDive to log fold change-based classification, as a function of increasing multicollinearity strength. The pancreas[5] dataset was downsampled to induce multicollinearity between females (F) and males (M) and non-diabetics (ND) and type 2 diabetics (T2D) (Cramer's V 0-1). Illustrated is an example peak with and without induced entanglement (Cramer's V 0.5) (F) and average precision across multicollinearity strength (Cramer's V 0-1) (G). **H)** Average biological conservation metric across the five datasets. Error bars represent the 95% confidence interval. **J)** Accuracy of predicting donor diabetes status using DeepDive versus baseline accuracy with XGBoost. **J-K)** Pearson $R^2$ of counterfactual, a baseline of acinar cells and beta cells from another donor compared to the hold-out donor beta cells (J) and selected example (K).

**Supplementary Figure 5:**
**A)** DeepDive basal embedding superimposed with cell type labels. B) Per peak DeepDive estimated effect sizes of BMI (x-axis) and diabetes status (y-axis). **C)** Enrichment of S-LDSC partitioned heritability in BMI and diabetes status associated peak sets for UKBB BMI, T2D, and handedness GWAS traits. **D)** Schematic of

XGBoost beta-cell state prediction. **E)** Train, validation, and test scores for training the XGBoost beta-cell state classifier. **F)** Alignment of beta-2 proportion across HbA1c level for author and XGBoost labels. **G)** Coefficients of gene expression and GSIS scores regression.
